# Decoding critical targets and signaling pathways in EBV-mediated diseases using large language models

**DOI:** 10.1101/2024.08.06.605212

**Authors:** Jingwen Yu, Yaohao Wang, Haidong Wang, Zhi Wei, Yonggang Pei

## Abstract

Epstein-Barr virus (EBV), a member of the gamma-herpesvirus, is the first identified human oncovirus and is associated with various malignancies. Understanding the intricate interactions between EBV antigens and cellular pathways is crucial to unravel the molecular mechanisms in EBV-mediated diseases. However, fully elucidating EBV-host interactions and the associated pathogenesis remains a significant challenge. In this study, we utilized LLMs to screen 36,105 EBV-relevant scientific publications, presenting a descriptive overview of the interactions between EBV antigens and host cellular pathways through comprehensive interaction networks. We described the critical roles of EBV antigens by constructing functional subsets of host proteins targeted by EBV antigens and illustrated the interactions using protein- protein interaction (PPI) networks. Furthermore, we developed an antigen-pathway network that highlights the connections with EBV-associated diseases, including DLBCL, BL, NPC, gastric cancer (GC), and post-transplant lymphoproliferative disorders (PTLD). Utilizing our dataset and public dataset, we validated the efficacy of using BLLF3-targeted TLR2-associated factors as marker genes for DLBCL. Next, we confirmed the co-expression of calcium pathway factors induced by LMP1 over- expression in BL. Finally, based on shared results suggesting that LMP1 actively regulates the glycolysis pathway, we identified and validated the correlation and co- expression of LMP1-induced PARP1, HIF1A, HOXB3, and key transcription-related factors, revealing the complete picture of LMP1’s influence on the glycolysis pathway. Our study presents a comprehensive functional encyclopedia of the interactions between EBV antigens and host signaling pathways across various EBV-associated disease contexts, providing valuable insights for the development of therapeutic strategies.

## Introduction

Epstein-Barr virus (EBV) was discovered in 1964 and has infected over 90% of the world’s population^1^. The oncovirus contributes to various human diseases, such as Burkitt lymphoma (BL), nasopharyngeal carcinoma (NPC), Hodgkin lymphoma (HL), diffuse large B cell lymphoma (DLBCL), multiple sclerosis (MS), and so on. In the 60 years since its discovery, the intricate roles of EBV in regulating interconnected host cellular pathways have provided valuable insights into the mechanisms underlying EBV-related disease progression. EBV infection in human cells triggers a series of changes in cellular biology including cell cycle transition, cell apoptosis, DNA mismatch repair, and multiple cellular signaling pathways. Such modulations are mediated by various EBV antigens, promoting viral latency and oncogenesis. For instance, EBV-encoded Latent Membrane Protein 1 (LMP1) mimics CD40 signaling via its two C-terminal activating regions, namely CTAR1 and CTAR2, which are shown to activate both the canonical and noncanonical NF-κB pathways^2–5^. EBV nuclear antigen 1 (EBNA1) inhibits the canonical NF-κB pathway by inhibiting the phosphorylation of IKKα/β^6^. Although the NF-κB pathway serves as a key factor in EBV latent infection and oncogenesis, the intricate and intertwined network of EBV antigens and NF-κB factors presents a significant challenge for the development of anti-EBV therapeutic strategies. Therefore, it is imperative to comprehensively examine the landscape of current studies on EBV antigens and their interactions with cellular signaling pathways to further explore the EBV-mediated pathogenesis.

Traditional literature mapping techniques, including systematic reviews and meta-analyses, often fall short of capturing the full breadth of EBV literature. However, the rapid development in machine learning offers promising tools for analyzing potential key factors in EBV-mediated pathogenesis from vast amounts of literature. A recent study characterizes the biomedical literature into a 2D map^7^. The embedding contains 21 million papers, presenting the unlimited possibilities of LLMs-assisted literature study. Another work introduced a knowledge graph of the m6A regulatory pathway utilizing GPT-4 to elucidate m6A’s regulatory mechanisms in cancer phenotypes across various cancers^8^. This burgeoning technology has catalyzed a proliferation of studies exploring its capabilities and applications in the fields of biological and medical research, driving innovation and prompting ongoing discourse on its potential impacts and future developments.

Our study aims to utilize LLMs for the first time to analyze decades of research on how EBV antigens regulate cellular factors and signaling pathways. We present a comprehensive framework of EBV antigens, their interacting host targets, and associated pathways, constructing an extensive network that delineates virus-host interaction. We then mapped the diverse background of EBV-related diseases for antigen-pathway relationships and revealed the critical interactions between EBV antigens and cellular pathways in EBV-mediated diseases. Collectively, this study offers a unique overview of current EBV-host interactions and provides valuable resources for the development of anti-EBV therapeutic strategies in the future.

## Methods

### Data sources

The data in this study were obtained from the PubMed database (https://pubmed.ncbi.nlm.nih.gov/) using EDirect. The initial data collection was performed on February 28, 2024. The final dataset comprises 36,105 EBV-related articles, including only original articles and reviews published in English. The RNA- seq dataset employed to investigate differentially expressed genes induced by LMP1 overexpression was obtained from the Gene Expression Omnibus (GEO) database under accession number GSE247495. Differential gene expression analysis was conducted using the R package DESeq2 (1.40.2). Genes with an adjusted p-value (padj) less than 0.05 and a log2 fold change greater than log2(2) were considered significantly differentially expressed.

### ChatGPT-4 prompt engineering

To ensure accuracy, we provided ChatGPT with the lists of EBV antigens and EBV-related diseases through system messages. The prompt for antigen-target- pathway summary is as follows: “If the abstract concludes that EBV-encoded proteins affect another protein or pathway, show me only the EBV protein, affected protein and pathway. If not, return ’Error’ and nothing else. Use a temperature of 0. All EBV proteins were listed before. The abstract: XXX”. The prompt for EBV-related disease mapping: “Return which specific EBV-related disease does the abstract attached in the end investigates. If unknown, return ’Error’ and nothing else. The abstract: XXX”.

To minimize costs, we initially utilized the free version GPT-3.5 Turbo via the OpenAI API. Items not resulting in “Error” were then processed using GPT-4. Next, we employed synonym substitution to standardize the naming of various aliases. For instance, “EBV nuclear antigen 1”, “EBNA1” and “EBNA-1” are all annotated as “EBNA1”; “EB2”, “Mta”, “SM” and “BS-MLF1” are all annotated as “BMLF1”. (Table S1)

### GPT-4 generated data evaluation

To evaluate the accuracy of the GPT-generated results, we randomly selected a total of 115 entries for manual inspection. The inspection focused on the EBV antigens mentioned in the corresponding articles, the host factors associated with these antigens, and the annotated KEGG pathways. The criteria for a “correct result” are as follows: (1) the article clearly describes the interaction between the EBV antigen and its corresponding host factors; (2) the article identifies the biological changes influenced by this interaction and their relevance to the annotated KEGG results.

### Annotation from GPT summarized pathways to KEGG database

The results generated by LLMs are highly variable. To establish a unique standard for readability, we annotated the LLM-generated pathway outputs to the KEGG pathway database due to its simplicity, accessibility, and widespread use within the academic community. This mapping was performed based on our customized R script and empirical manual substitution. A Sankey diagram shows a complete mapping of the substitution (Table S21).

### Gene enrichment analysis

LLM-acquired target host protein annotation is performed using the R package STRINGdb (2.12.1). Annotated IDs are submitted to UniProt for symbol conversion (https://www.uniprot.org/). The generated list of genes of interest (GOIs) was subjected to gene enrichment analysis using three primary tools: the KEGG database^9^, Reactome (https://reactome.org/), and STRINGdb.

### Omics analysis

PCA reduction and cohort analyses are performed using UCSC Xena^10^ (https://xena.ucsc.edu/) and GEPIA2 (http://gepia2.cancer-pku.cn/)^11^. The dataset comprised TCGA Diffuse Large B-Cell Lymphoma, GTEx Spleen and GTEx Blood. LMP1 overexpression RNA-seq data is normalized and analyzed using DESeq2.

### Visualization

The figures were created with BioRender (https://biorender.com), the R package wordcloud2 (0.2.1), circlize (0.4.16), or pheatmap (1.0.12) with customized scripts.

PPI network is visualized in Cytoscape (3.9.1) and colored based on protein family types. Other visualizations are made by a combination of R packages including ggplot2 (3.5.0), ggalluvial (0.12.5), and enrichplot (1.20.3).

## Results

### LLMs-assisted data mining reveals the key insights in EBV research

To create a literature pool for information extraction, our initial dataset comprised 36,105 EBV-related research articles available in the PubMed database as of the start of this project (February 28, 2024). These publications span a temporal range of 57 years, from 1968 to 2024 (Figure 1a). Overall, there has been a steady increase in the number of research articles related to EBV over these years, with an average of approximately 1000 publications annually.

**Figure 1.**
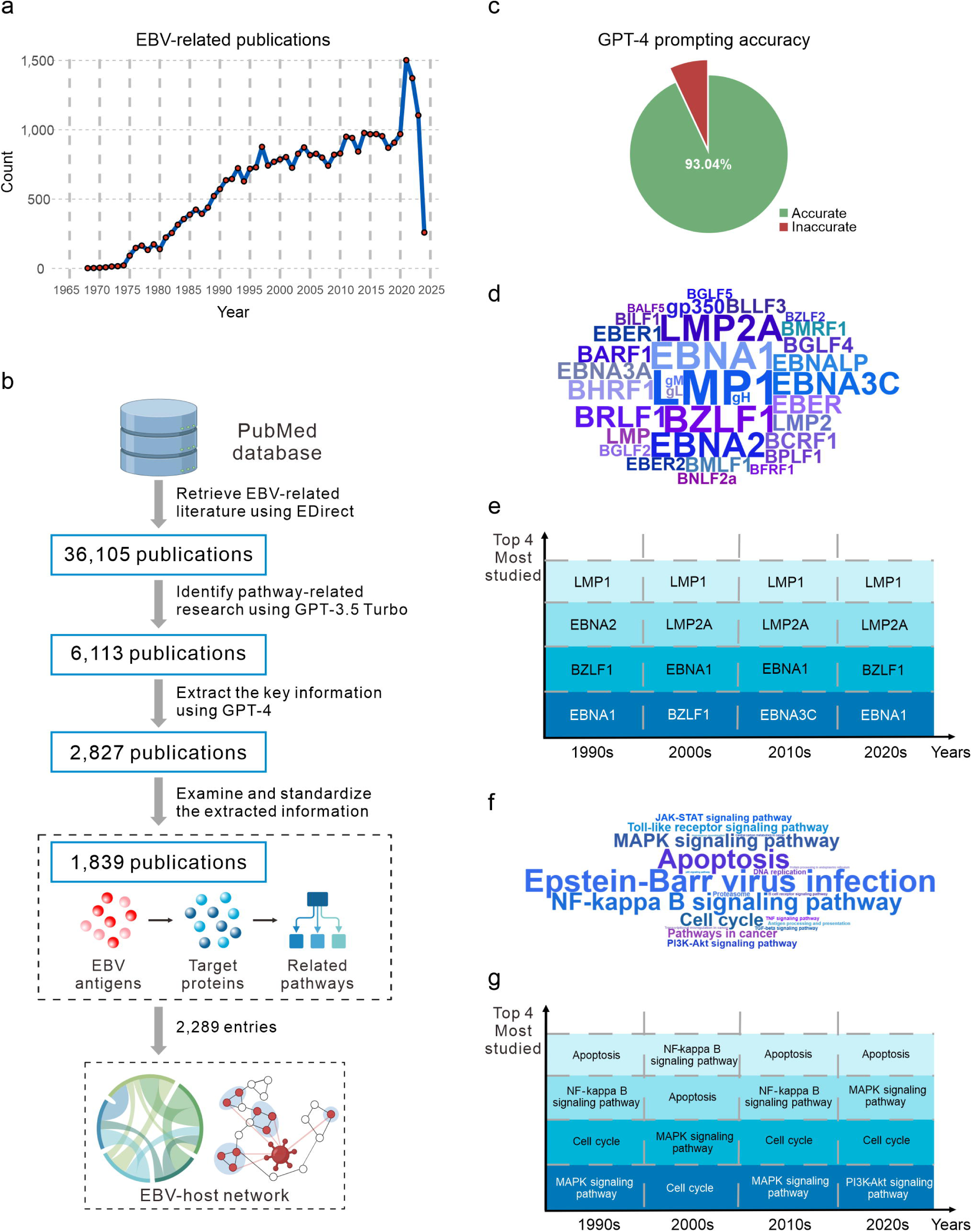
LLMs-assisted data mining reveals the key insights in EBV research. (**a**) The trend of EBV-related publications in different years. (**b**) The pipeline in this study is illustrated as a flowgraph. (**c**) Evaluation of GPT-generated output is conducted by examining 102 entries recorded for EBNA3C and LMP2. (**d**) The relative frequency of EBV antigens (log2 transformed) in our analysis is illustrated in the word cloud. (**e**) Statistical analysis highlights the research focus on various EBV antigens over time, showcasing only the four most studied antigens for each decade. (**f**) The relative frequency of EBV-related pathways was shown in the word cloud. (**g**) Statistical analysis highlights the research focus on various EBV-related pathways over time, showcasing only the four most pathways for each decade.

Next, to better characterize the current EBV studies, we summarized the available articles into a table of PubMed ID, EBV antigens, target proteins, and the related signaling pathways using ChatGPT-4. The returned dataset contained information from 2,827 publications related to EBV antigen-pathway interactions and was subsequently subjected to rigorous data reduction and filtering (Figure 1b). The final dataset includes 2,289 documented entries of EBV antigens and their associated pathways, derived from 1,839 publications (Figure 1b, Table S3). To validate the accuracy of the information extracted using ChatGPT, we manually compared a random subset of 115 entries (∼5%) of the GPT-4 generated summaries with the corresponding original abstracts. The results indicate that our prompts achieved an average accuracy of 93.04% (Figure 1c, Table S4), demonstrating that our GPT-4- based in-house pipeline can effectively analyze additional literature.

Analysis of EBV antigens and their impacts on cellular pathways reveals that LMP1 is the most frequently studied antigen, followed by EBNA1, BZLF1, LMP2A, and EBNA2, which are primarily latent or lytic proteins (Figure 1d). Throughout the 2010s, the significance of EBNA3C has increased, making it the fourth most studied antigen (Figure 1e). The most studied pathways focus on the mechanisms underlying EBV infection (Figure 1f). Pathways involving EBV-mediated cell apoptosis are the next most thoroughly studied areas. Abundant studies concentrate on the oncogenic mechanism of EBV infection, mainly on how EBV antigens regulate cell cycle and replication. The most extensively studied mechanisms include the NF-κB signaling pathway, MAPK signaling pathway, TLR signaling pathway, PI3K/Akt signaling pathway, TNF signaling pathway, and TGF-β signaling pathway (Figure 1f). On the temporal scale, research on cell apoptosis, NF-κB signaling pathway, cell cycle, and MAPK signaling pathway have always been the four unyielding topics (Figure 1g), suggesting their critical roles in EBV-mediated diseases.

### Pathway mapping reveals distinct signatures of EBV antigens

To clarify the critical pathways that EBV antigens regulate, we manually annotated the GPT-generated pathways to the KEGG pathway entries (Table S2). The chord diagram denotes the frequency of top 33 most studied EBV antigens and correlated pathways, indicating an intricate and interconnected network of cellular pathways induced by EBV infection (Figure 2a). To clearly illustrate the concentrated interactions between EBV antigens and their related pathways, we present a chord diagram as a heatmap (Figure 2b). In this figure, the pathways are clustered using the maximum linkage clustering method, and EBV antigens are arranged empirically based on their roles in EBV latent or lytic infection. We observed that EBV latent membrane proteins, LMP1 and LMP2A, exhibit similar interaction patterns across different host cellular pathways, although research on LMP1 has been more thorough as previously discussed (Figure 1d). The resemblance of these two EBV antigens lies in the shared role in their ability to reprogram host immune response through the MAPK, PI3K/Akt, or TNF pathways, and their oncogenic functions in regulating the p53 or PD1/PD-L1 checkpoint pathway. Despite the significant scarcity of LMP2A- related publications compared to LMP1, the studies collectively indicate that LMP2A contributes more to the BCR signaling pathway and the Wnt pathway than LMP1 (Figure 2b).

**Figure 2.**
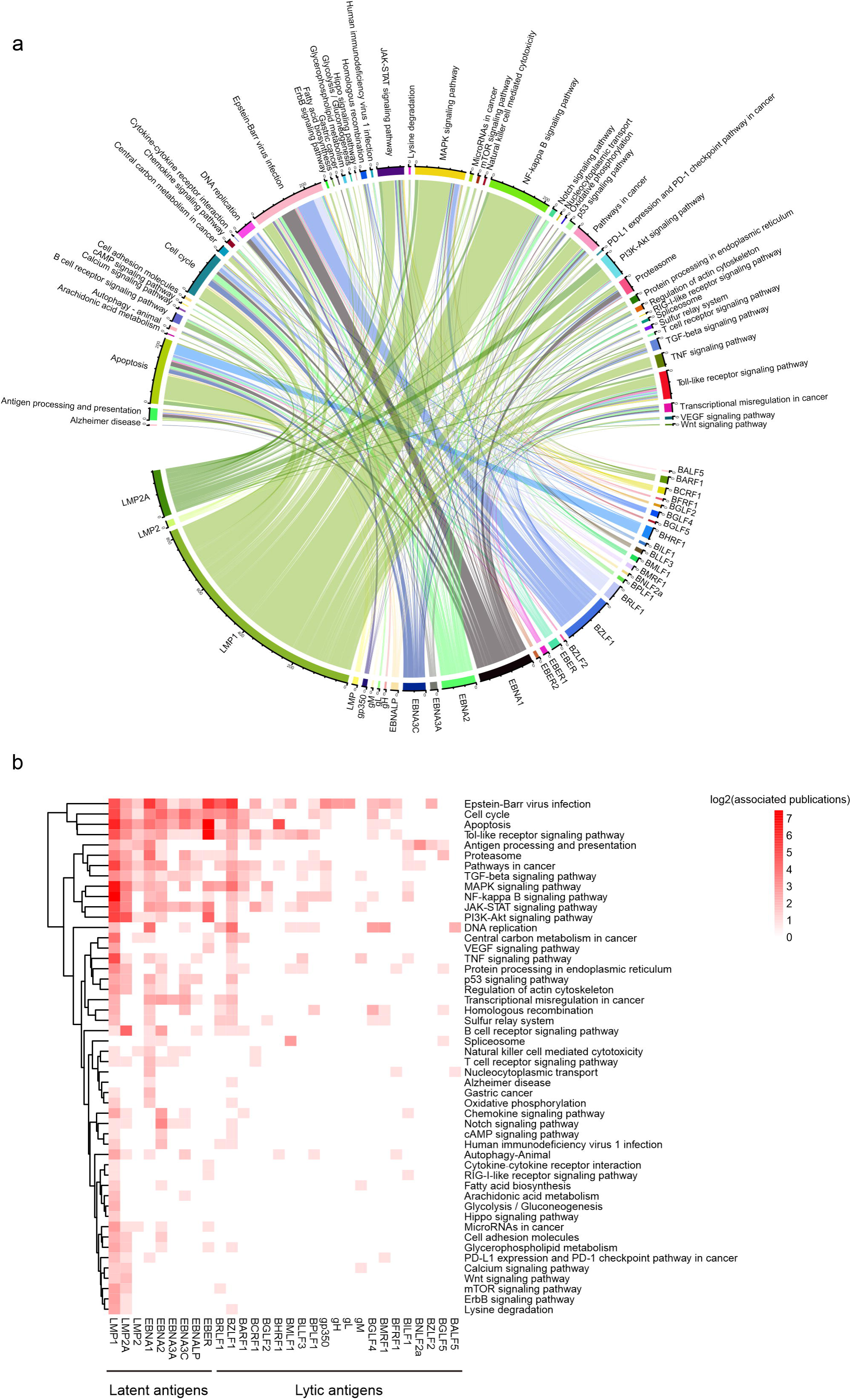
Pathway mapping reveals the distinct signatures of EBV antigens. (**a**) The chord diagram shows the linkage between the top 33 EBV antigens and their corresponding pathways. (**b**) Heatmap view of the linkage between EBV latent or lytic antigens and the associated pathways.

EBV nuclear antigens (EBNAs) contribute to the establishment and maintenance of viral latency, especially EBNA1, which is present in all EBV latent programs except for latency 0. EBNA1 directly binds to the host genome and is essential for the replication and maintenance of EBV episome^1^. It also serves as a transcription factor regulating multiple gene expression. Our analysis shows that EBNA1 differs from other EBNAs in terms of regulating DNA replication, and oxidative phosphorylation. EBNA2 plays a crucial role in the initial stage of EBV-mediated B-cell transformation. Interestingly, EBNA2 has been confirmed to bind to super-enhancer sites that are critical for the maintenance of EBV latency^12,13^. Our findings highlight the distinctive role of EBNA2 in cAMP signaling pathways, fatty acid biosynthesis, and cell adhesion lipid molecules, indicating its potential to facilitate B-cell transformation and proliferation through lipid metabolism. Similar to EBNA1, EBNA3A and EBNA3C are shown to interfere with the host genome and promote immune evasion, cell cycle control, and inhibition of apoptosis as transcription factors. EBNALP binds to EBNA2 and functions as a coactivator to promote the transformation of B-cells during EBV infection^1^. For immediate-early lytic genes that are responsible for EBV reactivation, both BZLF1 and BRLF1 are well-studied. BZLF1 activates lytic gene expression, initiating the lytic cycle, while BRLF1 synergizes with BZLF1 to promote lytic gene transcription, leading to viral replication and release. Studies on BZLF1 seem more diverse on multiple cellular oncogenic pathways.

Other EBV antigens have received relatively less attention compared to the previously mentioned latent and lytic antigens (Table S5). EBV glycoproteins play roles in infection and entry into host cells. Glycoprotein L (gL) cooperates with gH and gB to form the infection complex, mediating EBV-cell fusion and entry.

### Gene enrichment uncover defining characteristics of EBV-induced cellular modulations

To further investigate the associated diseases or pathways from the level of target proteins, we annotated the EBV research literature on a higher resolution after enrichment of GPT-generated target proteins using the KEGG database and the Reactome database. Disease-wise, KEGG enrichment revealed that EBV infection may be associated with traits typical of a wide range of diseases, including bladder cancer, pancreatic cancer, colorectal cancer, small cell lung cancer, and others (Figure 3a). The results suggest a possible correlation between these malignant diseases and EBV infection. However, it’s important to note that the KEGG database does not include EBV-related diseases like Burkitt lymphoma. Consequently, the diseases highlighted are primarily autoimmune-related and various forms of cancer (Figure 3a). The pathways enriched solely from target proteins and those retrieved from literature exhibit notable similarities, with extensive studies on pathways such as apoptosis, VEGF signaling pathway, TNF signaling pathway, Toll-like receptor signaling pathway, and IL-17 signaling pathway (Figure 3b). This indicates the coherence between GPT-generated pathways and the original articles. The last three pathways highlight the importance of studying how innate immunity is affected by viral antigens. The IL-17 signaling pathway triggers downstream NF-κB signaling pathway and is known to be a key pathogenic factor of autoimmune diseases^14^. However, the relationship between the IL-17 pathway and EBV infection remains unexplored. The high expression of IL-17 in multiple sclerosis patients could present a novel area of EBV research and provide druggable candidates for certain autoimmune diseases^15^.

**Figure 3.**
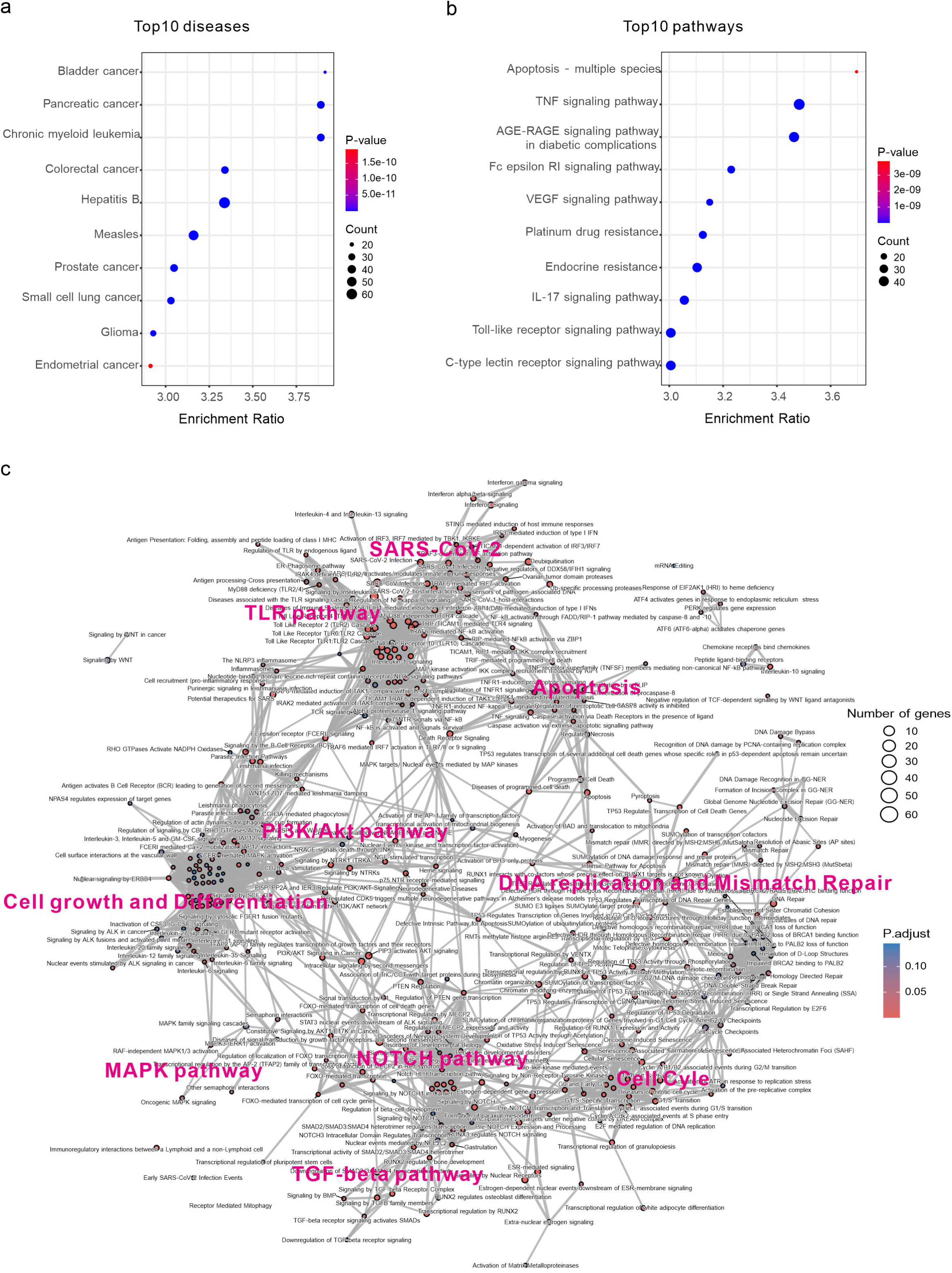
Gene enrichment and pathway analysis uncover defining characteristics of EBV-induced cellular modulations. (a-b) KEGG enrichment shows the top 10 EBV- related diseases or pathogens (a) and signaling pathways (b). (c) Reactome enrichment maps the landscape of EBV-targeted pathways and their correlations based on the shared host proteins.

To provide a more comprehensive overview of the functions of EBV-associated target proteins within their host, we used the Reactome database for annotation and identified 375 biological processes (p<0.05) related to EBV infection (Figure 3c).

This network shows that EBV-targeted host proteins predominantly form clusters in approximately 10 functional pathway groups. The largest clusters include innate immunity pathways, represented by the TNF signaling pathway, Toll-like receptor signaling cascades, and NF-κB signaling pathways. These are followed by cell growth and differentiation pathways, exemplified by the PI3K/AKT signaling pathways and related factors such as interleukins. Subsequently, DNA replication and mismatch repair, along with cell cycle-related pathways, are prominently featured, particularly those regulated by TP53. Moreover, the NOTCH signaling pathway and its regulation of the TGF-β signaling pathways mediated by the SMAD proteins are also critical for cell proliferation, differentiation, transitions, and apoptosis in our model of cellular signaling crosstalk^16^.

Intriguingly, we have identified some active mediators bridging the interconnected cellular signaling crosstalk (Figure 3c). These active members include deubiquitination and sumoylation that regulate protein modification and proteostasis; TP53-mediated cellular senescence on DNA replication and cell cycle; PTEN and downstream AKT signaling activation mediated by PIP3; BCR signaling for multiple pathways spanning the immune response to cell proliferation; Fc epsilon receptor (FCERI) signaling for the same function. Targets related to SARS-CoV-2 infection are enriched alongside the immune signaling pathways, suggesting a shared immune response elicited by EBV and SARS-CoV-2. This indicates the potential for developing broad-spectrum antivirals for both viruses by targeting shared oncogenic factors. Additionally, more discrete, smaller clusters were identified, including the Wnt signaling pathway, mRNA editing, as well as the unfolded protein response (UPR) essential for maintaining cellular homeostasis and survival under conditions of protein misfolding and ER stress.

### Analysis of host protein-protein interactions identifies key mediators in EBV- mediated diseases

To further identify critical cellular factors involved in EBV-associated diseases, we focused on EBV-targeted host proteins independent of pathway enrichment, which may provide insights into key target proteins. Bibliographic analysis of publications over time reveals trends in EBV-related host proteins (Figure 4a). p53 and c-Myc are the two most studied proteins across all periods. Genomic profiling has identified frequent TP53^17–19^ and MYC^20,21^ mutations or rearrangements in EBV-positive malignancies. CD23 and CD40 are two critical factors for EBV infection on the B-cell membrane. Earlier studies identified that EBNA2 specifically induces CD23 expression vital for B-cell transformation^22^ and verified the role of LMP1 in viral latency maintenance via mimicry of the CD40 signaling pathways^23^. As previously discussed in Figure 2, multiple EBV antigens including BHRF1 and LMP1 are associated with Bcl-2 related apoptosis in lymphoma and immortalized B-cells^19,24,25^.

**Figure 4.**
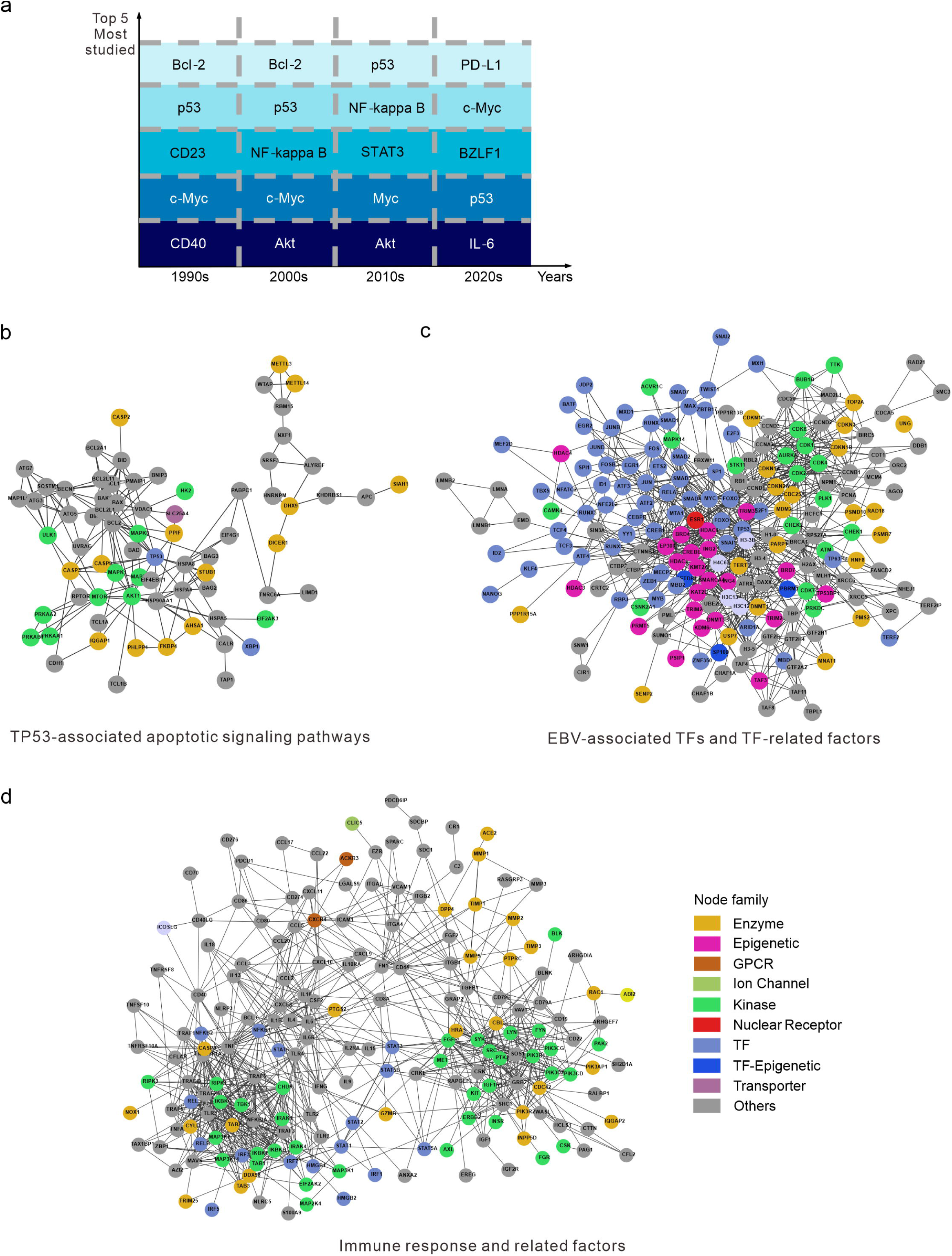
Analysis of host protein-protein interactions identifies key mediators in EBV-mediated diseases. (a) Statistical analysis reveals the focus of study every decade on EBV-targeted host proteins. TF, transcription factor; GPCR, G Protein- Coupled Receptor. (b-d) Top 3 PPI networks describe the basic functional clusters of EBV-targeted host proteins.

To better characterize the pathways and networks of EBV-related host factors described above, we annotated all the target proteins using the STRING database and performed protein-protein interaction (PPI) analysis. The top 3 enriched clusters are shown (Figure 4). The first PPI cluster centers around TP53 genes, apoptotic pathways, mTOR signaling, and the MAPK pathway cascades along with gene silencing mediated by DICER1 and m6A methylation (Figure 4b). This cluster reflects the role of EBV in suppressing the host apoptotic pathways and related signaling transduction pathways to promote viral latency and oncogenesis. Clinical trials and genomic profiling studies have identified the downregulation and mutation of TP53 in multiple EBV-associated cancers^26–29^. EBV-encoded microRNAs (EBERs) are shown to help maintain Burkitt’s lymphomas by targeting caspase 3 to inhibit apoptosis and aid the transformation of naïve B-cells^30^. Such inhibition usually works in harmony with other factors like Bcl-2. Besides triggering classic apoptotic pathways, EBV also bypasses the host antiviral systems by decreasing host Dicer protein via EBNA1- mediated microRNAs degradation^31^. Recent studies have also uncovered the role of circular RNAs in promoting the progression of EBV-associated gastric carcinoma through the transactivation of METTL3^32^. Along this pathway, caspases have been found to inhibit the m6A RNA modification process, thereby promoting EBV replication^33^. Additionally, EBV infection is known to induce TCL1 gene expression in Burkitt’s lymphoma, which also serves as an activator for the Akt pathway^34^.

The second cluster highlights EBV’s roles in transcriptional regulation and epigenetic modification, particularly through transcriptional factors (TFs) (Figure 4c). MYC is a key TF associated with multiple cancers including EBV-associated lymphomas^35^. Moreover, MYC abundance represses EBV reactivation and controls the lytic switch in Burkitt lymphoma^36^. 3D genome mapping of EBV-positive lymphoblastoid cell lines (LCLs) revealed distinctive EBV super-enhancer (SE) targets like MCL1, IRF4, and EBF^37^. Bioinformatic analysis identified that H3K27ac SEs overlap with STAT5 and NFAT targets, indicating their roles in promoting EBV latency^13^. On a larger scale, broad screening of TFs integrated over 700 high- throughput sequencing datasets and provides an atlas of EBV-related transcriptomic and epigenetic host-virus regulatory interactions^38^. These EBV-specific TFs, such as SPI1, EBF1, and PAX5, collectively contribute to EBV-induced oncogenesis and latency^13^.

The last cluster primarily comprises factors related to the innate immune response, including cellular receptors, cytokines, and chemokines that regulate immune cell activation, inflammatory response, and the tumor microenvironment (TME) (Figure 4d). EBV orchestrates immune evasion by cellular receptors like CD40, CD80, IL4, IL6, IL10 and their ligands^39,40^. For example, LMP1 mimics CD40 to activate downstream NF-κB pathways to induce B-cell proliferation^23^. After infection, EBV hijacks host signaling transduction factors like STATs^41,42^ and IRFs^43^ to promote cell survival and proliferation. LMP1 and LMP2A then activate the JAK/STAT pathway, to facilitate cell growth factors and anti-apoptotic signaling. Meanwhile, EBV and other herpesviruses induce the PI3K/Akt pathway to aid viral infection, latency, and reactivation^44^. This cluster is also rich in enzymes, multiple cellular growth factors (CGFs), and extracellular matrix (ECM) proteins, which represent the invasiveness of EBV-infected cells. Early research discovered that viral LMP1 and BZLF1 proteins up-regulate the expression and activity of matrix metalloproteinase 1 (MMP1), and thereby confer the invasive properties of the cells^45^. Besides common growth factors, we also identified Angiotensin-Converting Enzyme 2 (ACE2) in this cluster. Study indicates that the lytic replication of EBV induces ACE2 expression in human epithelial cells, which enhances the entry of SARS-CoV-2^46^.

### Exploring the critical relationships in EBV-mediated diseases

Due to the complex background of EBV pathogenesis, we next seek to elucidate the influence of EBV antigens on host pathways in the context of multiple EBV-associated malignancies (Figure 5a-b). In this section we demonstrate how our dataset and generated PPI networks can be utilized to assist EBV-host interaction studies using basic omics analyses. We plot the connections identified for EBV antigens and host pathways onto circular connection hubs for DLBCL, BL, NPC, GC, and PTLD (Figure 5c, Figure 6a and Figure S1).

**Figure 5.**
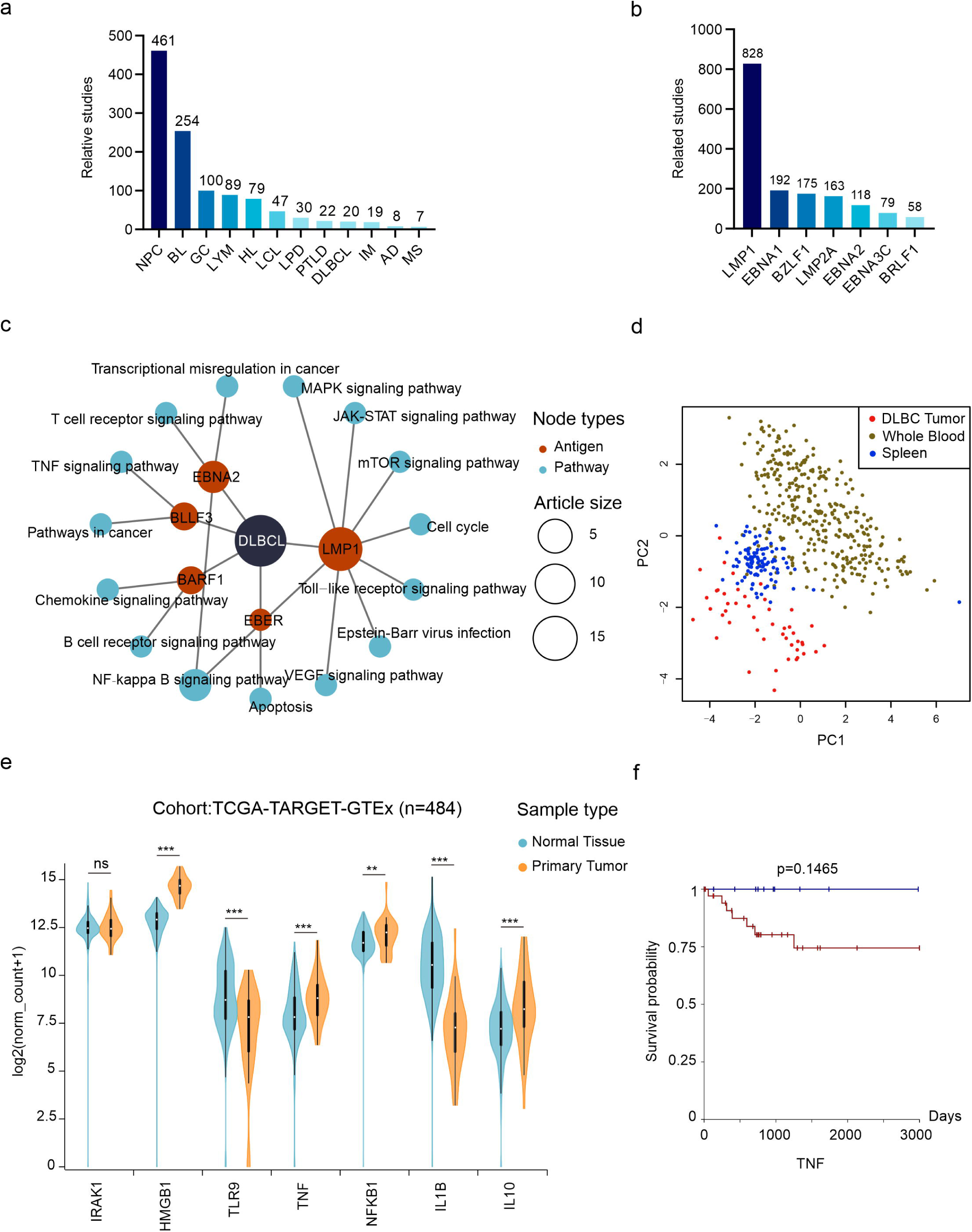
Revealing the key relationships in EBV-mediated DLBCLs. (a) The bar plot shows the numbers of total articles of the top 12 most studied EBV-related diseases. (b) The bar plot shows the numbers of total articles of the top 7 most studied EBV antigens. (c) The circular hub graph elucidates the DLBCL-related antigen-pathway relations. (d) PCA analysis highlights distinct DLBCL tumor samples using TLR2- related marker gene expressions. (e) The TLR2-related marker gene expressions between normal tissue and tumor tissue. (f) Survival plot shows high TNF expression leads to less survival probability (red indicates samples with normalized TNF counts > 7.907 and blue indicates less).

**Figure 6.**
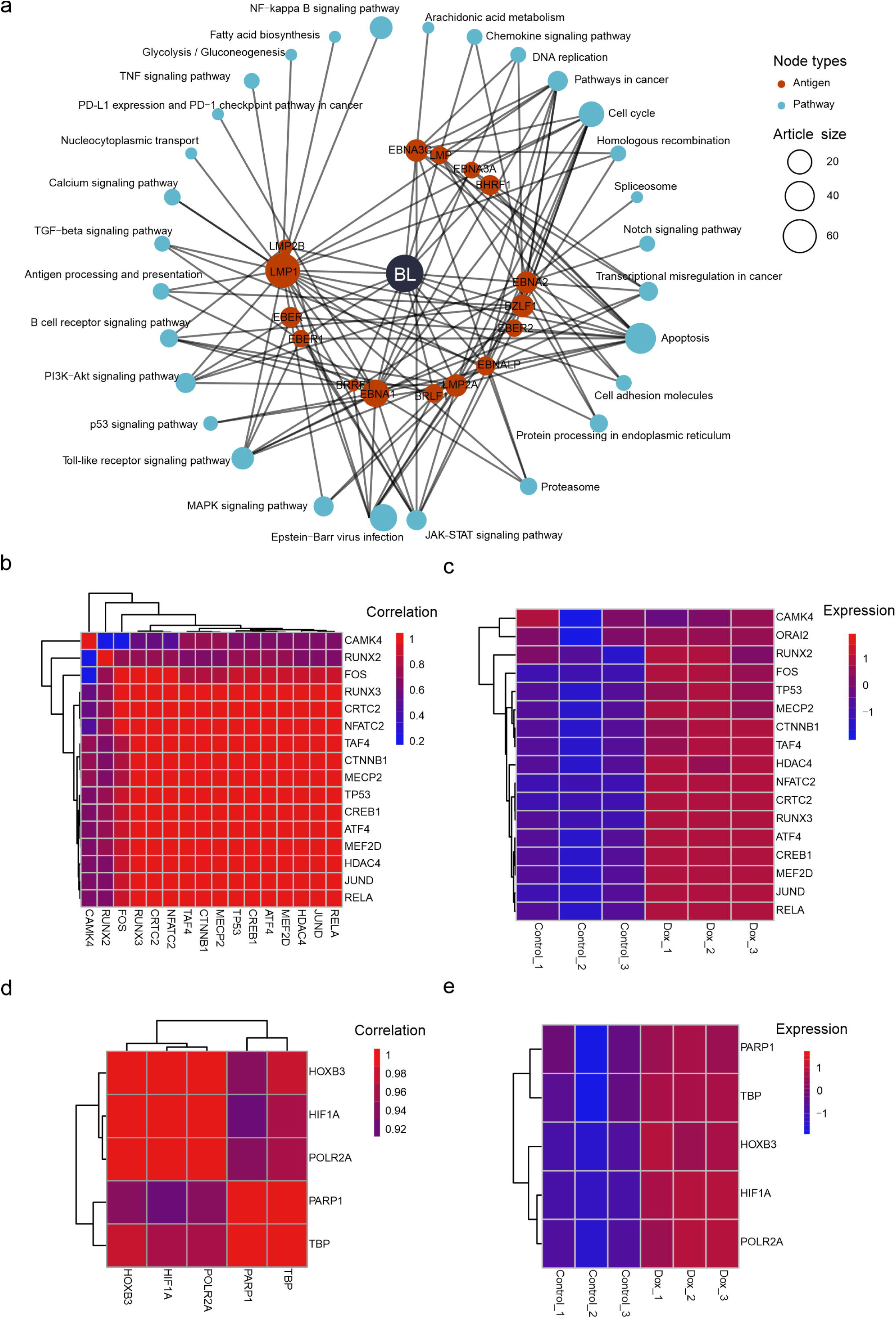
Exploring the functions of EBV antigens in BL. (a) The circular hub graph elucidates the BL-related antigen-pathway relations. (b-c) The CAMK4-related factors are co-expressed by LMP1 overexpression. (d-e) The PARP1-related factors are co-expressed by LMP1 overexpression.

Approximately 9% of DLBCLs are EBV positive with type II lantency^1^. In DLBCL-centered hubs, we noticed the special linkage of EBV encoded dUTPase BLLF3 and the TNF signaling pathways (Figure 5c). This indicates that BLLF3 induce DLBCL through the TNF pathways. dUTPase is crucial for maintaining the balance of intracellular nucleotides, particularly during DNA replication^47^. Our dataset pointed to study exploring BLLF3 and its interaction with host factor TLR2^48^. Our PPI networks show several known key factors closely interacting with TLR2 in the background of EBV infection, such as IRAK1, HMGB1, TLR9, TNF, NFKB1, IL1B and IL10. Next, we conducted pan-cancer analysis to examine the utility of these proteins in distinguishing between DLBCL tumor and normal samples (Figure 5d). The results indicated clear difference between the tumor and samples from normal tissues, suggesting their potential as marker genes for DLBCL diagnosis. As DLBCL does not have corresponding gene expression data for normal tissue in TCGA, we used whole blood and spleen, both of which are immune cells, as normal samples for comparison. Taken apart, we noticed the high expression of HMGB1, TNF, NFKB1 and IL10 in DLBCL tumor samples which function in TNF pathway (Figure 5e). At last, we examined the correlation of TNF expression and DLBCL patients’ survival probability (Figure 5f). The results show the survival possibility of patients with high TNF expression dropped to 75% in approximately four years while samples with low expression survived 100%. These findings above show the potential in studying BLLF3 mediated TNF pathway activation in the carcinogenesis of DLBCL.

LMP1 is the most studied EBV antigens in our literature pool (Figure 5b). In BL, literature indicates that LMP1 activates the calcium signaling pathway by overexpressing the host factor CAMK4^49^ (Figure 6a, Table S3). Our PPI network shows that HDAC4 and CREB1 directly interact with CAMK4, and the secondary factors that interact with these two, such as MEF2D, RUNX3, RUNX2, NFATC2, JUND, FOS, RELA, TP53, TAF4, MECP2, CTNNB1, ATF4, and CRTC2 (Figure 4c).

We observed that although a dox-induced overexpression of LMP1 in the Akata cells did not result in a significant increase in CAMK4, its interacting proteins listed above exhibited a co-expression pattern and consistent upregulation (Figure 6b, c). This suggests that the calcium signaling pathway may play a role in the oncogenesis of BL by a LMP1-dependent manner.

Finally, we locate studies working on similar pathways associated with the same antigens across cell lines from different EBV-related malignancies to compensate the utility. For instance, studies on BL and NPC both demonstrated that LMP1 is associated with glycolysis (Figure 6a, Figure S1). EBV typically establishes type I latency in BL. Overexpressing LMP1 in BL cell lines suggests that LMP1 activates PARP1 to induce glycolysis and promote oncogenesis in a HIF-1 α -dependent manner^45^. Study in NPC cell lines demonstrates that LMP1 represses Hox genes via stalling the key mediators of DNA transcription like RNA pol II to modulate glycolysis^51^. These studies in cell lines with different EBV latencies demonstrate the role of LMP1 in regulating glycolysis. We next explore the transcriptomic correlation for PARP1, HIF1A, HOXB3, and key transcription-related factors (Figure 6d-e). Result show high correlation, which suggests a more complete mechanism that LMP1 activates PARP1, which subsequently binds to HIF-1α and RNA pol II and activates HoxB3, finally modulate glycolysis.

## Discussion

This study, for the first time, presents a mapping of EBV antigens, host factors, and cellular pathways from available EBV-related articles using LLMs. We have summarized the major pathways associated to each EBV antigens, and provided a descriptive overview of core PPI clusters. At last, we provided examples of ustilizing our dataset for marker gene analyses.

However, certain limitations in our research need to be addressed. While our initial data pool contained 36,105 EBV-related publications, our strict prompts and data filtering reduced this to 2,289 entries, encompassing 1,839 unique publications (∼5.1%) for downstream analyses. If loosening the criteria, we will be able to obtain more entries for deeper analyses, although the accuracy may decrease. Secondly, our literature study was based solely on conclusions drawn from abstracts of the available publications which has its intrinsic limitation in the study of network biology. Although we used the frequency of topics as a proxy for their importance in the EBV antigen-pathway mapping, the sheer number of publications may not fully represent actual biological significance. Occasionally, a higher citation rate may influence their proportions, potentially translating non-experimental results. Future efforts may focus on using supervised machine learning algorithms to explore the transcriptomic networks of EBV-induced cellular changes for enhanced utility and usability. In the last part, we showcase through basic omics analyses that our EBV literature analysis bears the potential to inspire researchers to engage with former published work of scientists worldwide for novel effort in illustrating the mechanism of EBV-host interaction. Finally, EBV latency types are critical in understanding EBV gene expression and virus-host interaction. However, such information is largely unattainable in abstracts. Future research based on more powerful tools can extract the main text for related information.

This study represents a pioneering effort in the application of LLMs in EBV- related research, highlighting the potential of LLMs in advancing our understanding of viral interactions and cellular pathways. Unlike broad-scale context generation, our study clarifies the relationships between EBV antigens and various cellular pathways based on the manual annotation of GPT-generated summaries, presenting networks of EBV-related host proteins for future research. Beyond EBV, the big data era and advancements in various LLMs underscore the urgent need for advanced, digitized platforms in all areas of viral research. Such tools are pivotal for accelerating research progress, advancing basic virology, and fostering the development of novel antivirals.

## Acknowledgments

We thank all members of our laboratory for helpful advice and assistance.

## Author contributions

Y.P. conceived the application of LLM in the EBV-related literature analysis. J.Y. designed and carried out the data mining and visualization. Y.W., H.W., and Z.W. provided expertise in technical support and data analysis. J.Y. wrote the manuscript and Y.P. revised the manuscript.

## Supplementary materials

Figure S1: Determining the key relationships in EBV-mediated NPC, PTLD, and GC. (a-c) The circular hub graph elucidates the NPC-related, PTLD-related and GC- related antigen-pathway relations. NPC, nasopharyngeal carcinoma; PTLD, Posttransplant lymphoproliferative disorders; GC, gastric cancer.;

Table S1: The formal names and aliases used in GPT-generated term substitution

Table S2: The correspondence between GPT-4 generated pathways and manually annotated pathway to KEGG entries;

Table S3: The final dataset includes 2,289 entries of EBV antigens and their associated pathways;

Table S4: Validation for GPT-generated results;

Table S5: Ranking of most studied EBV antigens (with >10 related articles);

