## Supplementary figures and images for "Decoding critical targets and signaling pathways in EBV-mediated diseases using large language models"

### Figure S1

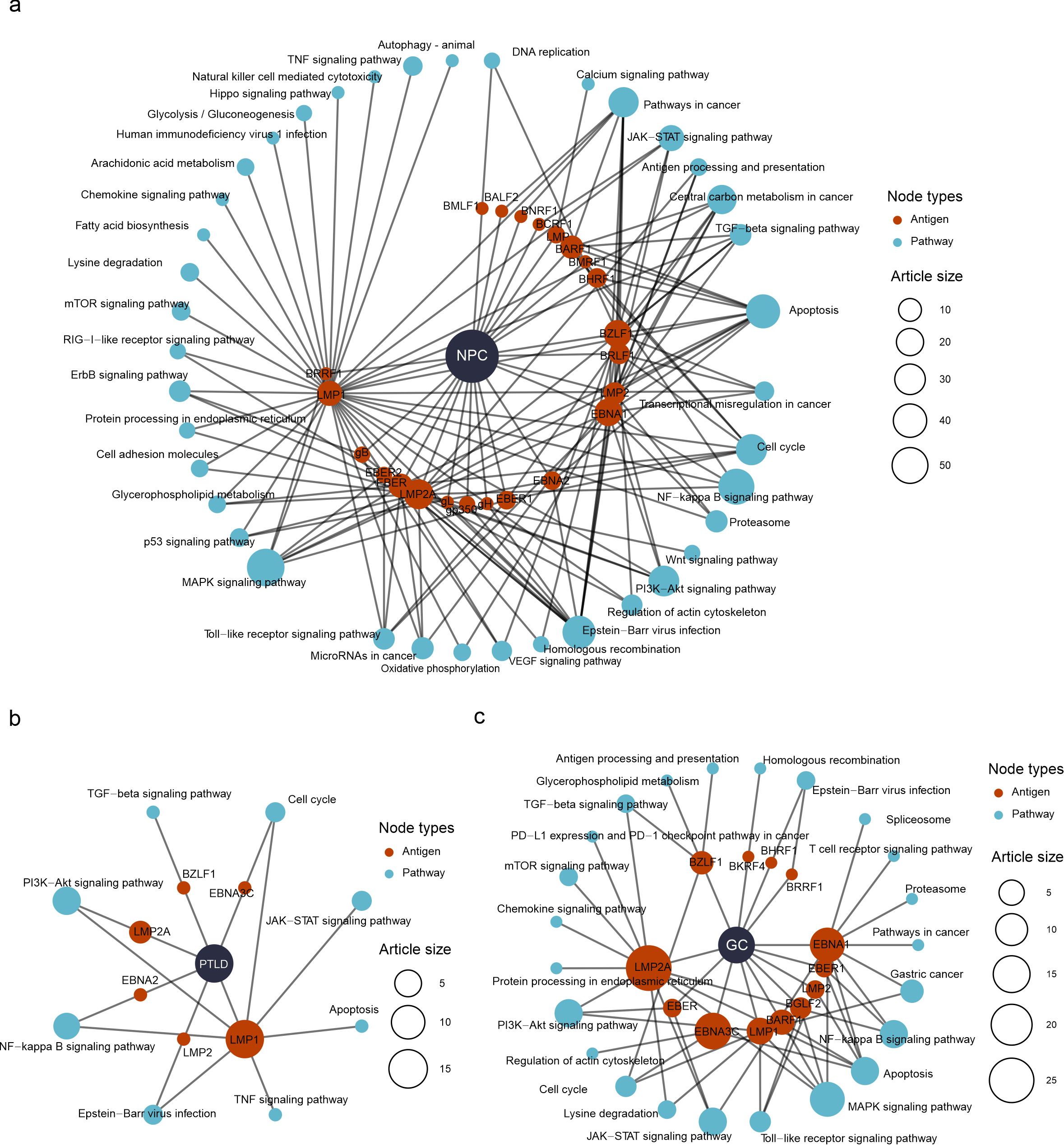
